# A Lysozyme Murein Hydrolase with Broad-spectrum Antibacterial Activity from *Enterobacter* Phage myPSH1140

**DOI:** 10.1101/2022.04.06.487332

**Authors:** Nachimuthu Ramesh, Prasanth Manohar, Kandasamy Eniyan, Loganathan Archana, Sudarsanan Athira, Belinda Loh, Long Ma, Sebastian Leptihn

**Affiliations:** Antibiotic resistance and Phage therapy Laboratory, School of Biosciences and Technology, Vellore Institute of Technology, Vellore, India; Zhejiang University-University of Edinburgh (ZJU-UoE) Institute, Zhejiang University, Haining, Zhejiang 314400, P.R. China; Fraunhofer Institute for Cell Therapy and Immunology (IZI), Department of Antimicrobial Biotechnology, Leipzig, Germany; Department of Infectious Diseases, Sir Run Run Shaw Hospital, Zhejiang University School of Medicine, Hangzhou, P.R. China; University of Edinburgh Medical School, Biomedical Sciences, College of Medicine &Veterinary Medicine, The University of Edinburgh, 1 George Square, Edinburgh, EH8 9JZ, United Kingdom

**Keywords:** Endolysins, Peptidoglycan hydrolase, Gram-negative pathogens, Phage therapy, Antimicrobial Proteins

## Abstract

Bacteriophages and bacteriophage-derived peptidoglycan hydrolases (endolysins) present promising alternatives for the treatment of infections caused by multi-drug resistant Gram-negative and Gram-positive pathogens. In this study, Gp105, a putative lysozyme murein hydrolase from *Enterobacter* phage myPSH1140 was characterized *in silico, in vitro* as well as *in vivo* using the purified protein. Gp105 contains a T4-type lysozyme-like domain (IPR001165) and belongs to Glycoside hydrolase family 24 (IPR002196). The putative endolysin indeed had strong antibacterial activity against Gram-negative pathogens including *E. cloacae*, *K. pneumoniae*, *P. aeruginosa*, *S. marcescens*, *Citrobacter* sp. and *A. baumannii*. Also, an *in vitro* peptidoglycan hydrolysis assay showed strong activity against purified peptidoglycans. This study demonstrates the potential of Gp105 to be used as an antibacterial protein to combat Gram-negative pathogens.

## Introduction

Antibiotic resistance is a global health issue of major concern. Because of the rapid spread of resistance, new antimicrobial agents are needed and those showing both, broad spectrum effects and low levels of resistance, would be beneficial (1). Gram-negative bacteria (GNB) are known to cause the majority of serious infections in humans (2,3). Causative agents that predominantly belong to the *Enterobacteriaceae* family include *Escherichia coli, Klebsiella pneumoniae, Enterobacter cloacae*, but also other non-fermenting GNB such as *Pseudomonas aeruginosa* and *Acinetobacter baumannii*. These bacterial pathogens have been declared a priority by the WHO, with the Disease Society of America terming them “ESKAPE” pathogens comprising *Enterococcus faecium, Staphylococcus aureus, Klebsiella pneumoniae, Acinetobacter baumannii, Pseudomonas aeruginosa* and *Enterobacter* species. These bacteria are known to cause the most critical infections in humans and the development of treatment possibilities are urgent (4–6). *Enterobacter* species include *E. cloacae, E. hormaechei, and E. asburiae* some of which cause infections of the urinary tract but also bloodstream infections (7,8). Globally, the infections caused by antibiotic-resistant *Enterobacter* are concerning; therefore, antibiotic alternatives are urgently needed.

Phage therapy may be an effective solution to overcome the antibiotic crisis. This is due to the availability, selective killing ability, synergy with antibiotics and low toxicity, among others (9–12). However, using “living viruses” may not be accepted by the general population as the idea might cause unfounded fears and rejection of such treatment. Phages produce bactericidal proteins which might be a good alternative to both, phages and chemical antibiotics. Endolysins are phage-derived enzymes that, during lysis, aid in the release of phage particles from the bacterial host by disintegrating the peptidoglycan (PG) component of the bacterial cell wall (13–15). Endolysins are amongst the most promising phage-derived proteins candidates to fight bacterial infections. While endolysins work by breaking down peptidoglycan of the host cell wall from within, the enzyme can also act from the outside, with studies demonstrating their rapid antibacterial activity. Another advantage is that endolysins have a broader host range compared to phages, while still exhibiting a narrow activity spectrum compared to chemical antibiotics thus leaving the microflora intact (16–19). Highly promising is also the observation that no resistance to endolysins has been reported thus far (16). Over the past few years, endolysins have received wide-ranging attention as therapeutic agents for the clinic (17–19). Several clinical studies have demonstrated the efficacy of endolysins to eliminate bacterial infections caused by multidrug resistant pathogens (20,21).

*Enterobacter* phage myPSH1140 was isolated against a clinical strain of *E. cloacae*, which was reported to infect four different species of *Enterobacter* (22). The genome size was found to be 172,614 bp encoding 102 functional proteins and 138 hypothetical proteins (NCBI accession number: MG999954). In this study, a lysozyme murein hydrolase (Gp105) has been identified in the phage and was first characterized *in silico*, revealing the protein to contain a T4-type_lysozyme domain and that belongs to Glycosyl hydrolase family 24. This protein family is so far uncharacterized in bacteriophages infecting *Enterobacter* species. In order to evaluate the potential of the protein to be used as a therapeutic, the coding gene was cloned and overexpressed in *E. coli*. Subsequent tests using the purified protein showed that Gp105 is able to hydrolyse purified peptidoglycan *in vitro* and exhibits strong antibacterial properties against a wide range of pathogens including *E. cloacae*, *K. pneumoniae, P. aeruginosa, S. marcescens, Citrobacter* sp. and *A. baumannii*.

## Materials and Methods

### In silico analysis of Gp105

Nucleotide and protein sequences that were analyzed by *in silico* methods were retrieved from NCBI database (http://www.ncbi.nlm.nih.gov). The multiple sequence alignment was carried out using Clustal Omega Program (https://www.ebi.ac.uk/Tools/msa/clustalo/) and the alignment file was used to generate phylogenetic relationship using MEGA 7.0 software (23). Signal peptides and sub-cellular localization was predicted by SignalP (http://www.cbs.dtu.dk/services/SignalP/) and TMHMM (http://www.cbs.dtu.dk/services/TMHMM-2.0/) program, respectively. The secondary structure of the protein was predicted using JPred Program (http://www.compbio.dundee.ac.uk/jpred/). The tertiary structure of the protein was established using SWISS-MODEL server (http://swissmodel.expasy.org/) based on homology modeling and analysed by Pymol (https://pymol.org/2/).

### Bacterial strains, plasmids and chemicals

*Escherichia coli* DH5α and *Escherichia coli* BL21 (DE3) were used for cloning and expression studies, respectively. The clinical strains of *Escherichia coli*, *Klebsiella pneumoniae*, *Enterobacter cloacae*, *Acinetobacter baumannii*, *Salmonella* typhi, *Pseudomonas aeruginosa, Serratia marcescens, Citrobacter* sp., *Enterococcus* sp. and *Staphylococcus aureus* were used from the Antibiotic Resistance and Phage Therapy Laboratory, VIT, Vellore. T4 DNA ligase and restriction enzymes BamHI and XhoI were obtained from New England Biolabs (USA). Oligonucleotide primers were synthesized by Eurofins Scientific India Pvt. Ltd (India). The plasmid-pET28a was purchased from Novagen (Germany). Ni-NTA agarose was purchased from G-Biosciences (USA), and plasmid isolation and gel extraction kits were obtained from Takara Biotech (USA). All analytical grade chemicals and antibiotics were purchased from Himedia chemicals (India). Standard recombinant DNA techniques were used as described elsewhere (24).

### Phage propagation and DNA extraction

Phage enrichment method was used to propagate phage myPSH1140 against the host bacteria, *E. cloacae*. Briefly, 1 mL of phage lysate was added to 3 mL of overnight grown bacterial cultures (host bacteria), and the mixture was incubated at 37°C for 24 hours in a shaking incubator (150 rpm). The mixture was centrifuged at 6000 x g for 15 min and the supernatant was filtered through a 0.22-micron syringe filter. The filtrate was tested for phage activity using spot test and double agar overlay method as previously described (22).

Phage DNA was extracted from the enriched phage lysate using phenol-chloroform (24:1) method. Briefly, the phage particles were treated with DNase (20 mg/mL) and RNase (0.5 mg/mL) for 1 h at 37°C to remove any host DNA and RNA. To the mixture, 20% PEG and 1.6 M NaCl was added and stored at 4°C for 1 h. The content was centrifuged at 12,000 x g for 10 min and to the pellet, phage lysis buffer (50 μL of 10% SDA, 50 μL of 0.5M EDTA, and 5 μL of 10 mg/mL proteinase K) was added, mixed well and incubated overnight at 50°C. Equal volume of phenol: chloroform: isoamyl alcohol (25:24:1) was added and centrifuged at 12,000 x g for 15 min (repeated twice). The aqueous phase was extracted into a new tube and an equal volume of 100% isopropanol was added to precipitate the DNA at −20°C for 6 hours. The precipitated DNA was pelleted, washed with 100% ethanol and air dried. The purified phage DNA was visualized on 0.8% agarose gels.

### PCR Amplification

The nucleotide sequence of gp105 gene was retrieved from NCBI (http://www.ncbi.nlm.nih.gov) for designing gene specific primers. PCR was performed using the genomic DNA of *Enterobacter* phage myPSH1140 as template. The gene was amplified using gene-specific primers listed in Table 1, containing the BamHI (Forward primer) and XhoI (Reverse primer) restriction sites. The gene was amplified under the following conditions: 95°C for 5 min, 25 cycles (95°C 1 min; 55°C 1 min; 72°C 1min), and final extension of 72°C for 10 min. The PCR products were analyzed on a 1.2% agarose gel.

**Table 1.**
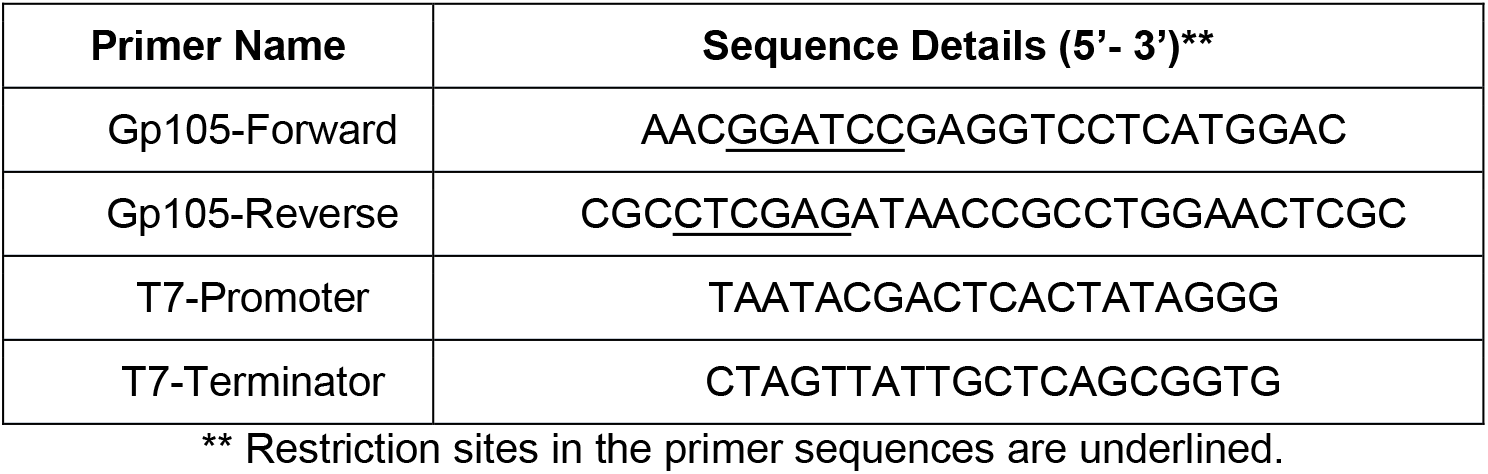
The primer sequences used to amplify the gene, Gp105.

### Cloning and expression

The PCR products generated was gel purified, digested and ligated into pET28a vector. The ligation mixture was transformed into *E. coli* DH5α cells and the colonies obtained were screened using T7 Promoter and T7 Terminator primers (Table 1). Recombinant plasmid was isolated from the positive colonies using plasmid miniprep kit and verified by DNA sequencing.

The recombinant plasmid pET28a-gp105 was transformed into *E. coli* BL21 (DE3) cells for expression studies. A single colony of *E. coli* BL21 (DE3) containing pET28a-gp105 was inoculated in 5 mL of LB medium containing kanamycin (50 μg/mL) and grown at 37°C overnight. A 1 mL of this culture was added in 100 mL fresh LB medium containing kanamycin (50 μg/mL) and incubated at 37°C to reach the log phase. To the growing culture of *E. coli* BL21 (DE3) containing pET28a-gp105, IPTG was added to a final concentration of 1 mM and incubated at 37°C for 3 hours. After induction, cells were harvested by centrifugation at 8,000 × g for 10 min and the total protein was analyzed (both the control and induced cells) in 12% SDS-PAGE. The proteins were visualized by staining the gel with Coomassie blue (0.05%) for 30 min in a rocking shaker.

### Protein purification

*E. coli* BL21 (DE3) cells transformed with pET28a-Gp105 were grown at 37°C in LB broth containing kanamycin (50 μg/mL) until log phase and the expression was induced by the addition of 1 mM IPTG for 3 hours at 37°C. The cells from the 100 mL culture were resuspended in 5 mL of lysis buffer (50 mM phosphate buffer, pH 8.0, 300 mM NaCl, 5 mM imidazole, 1 mM PMSF and 10 μg/mL lysozyme). After one cycle of freeze-thaw, the cells were lysed by sonication (30s pulse x 30s pause). The soluble fraction was separated by centrifugation at 15,000 × g for 30 min at 4°C and loaded on to a pre-2 mL (bed volume) Ni–NTA agarose column. After washing the column with 30 bed volumes of Wash buffer A (50 mM Phosphate Buffer, pH 8.0, 300 mM NaCl, 20 mM imidazole) and Wash buffer B (50 mM Phosphate Buffer, pH 8.0, 300 mM NaCl, 40 mM imidazole) at a flow rate of 1.0 mL/min, the bound protein was eluted in Elution buffer (50 mM phosphate buffer, pH 8.0, 300 mM NaCl, 200 mM imidazole). The purified proteins were quantified using Bradford method.

### Peptidoglycan assay

#### Peptidoglycan purification

The bacterial peptidoglycan (PG) from *Enterobacter cloacae* and *Staphylococcus aureus* was obtained as described earlier by Santin et al. (25). Briefly, 400 mL of bacterial cells at OD_600_ 1-1.2 was harvested by centrifuging at 10,000 x g for 20 min at 4°C. The bacterial pellet was resuspended in 20 mL of buffer I (20 mM Tris-Cl and 100 mM NaCl, pH 8). The bacterial cells were broken by passing (3-5 times) through a syringe needle (0.4-by 20 mm needles) and the cell wall was collected by ultracentrifugation at 90,000 × g for 45 min at 4°C (Beckman Coulter, Inc.). To the pellet, 10 mL of 8% SDS was added and incubated at 96°C for 1 h. The peptidoglycan fraction was collected by ultracentrifugation at 90,000 × g for 45 min at 25°C and the obtained peptidoglycan was resuspended in 10 mL of 0.5 M NaCl and 10 mL of 8% SDS. The peptidoglycan was mixed thoroughly and incubated at 96°C for 30 min. The peptidoglycan was then pelleted by ultracentrifugation at 90,000 × g for 30 min at 25°C and resuspended (twice) in 10 mL of sterile distilled water. Then, the peptidoglycan was resuspended in 10 mL of buffer II (20 mM Tris-HCl, pH 7.2, 50 mM NaCl supplemented with 200 μg/mL α-amylase and 200 μg/mL pronase), and incubated at 37°C for overnight. After incubation, 10 mL of 8% SDS was added and incubate for additional 1 h at 96°C. The peptidoglycan was pelleted by ultracentrifugation at 90,000 × g at 25°C for 30 min and the obtained peptidoglycan fraction was resuspended (twice) in 10 mL of water. The pellet obtained in final step was resuspended in 1 mL of water. The peptidoglycan was stored at 4°C until use.

#### Peptidoglycan hydrolysis assay

The hydrolysis of peptidoglycan at different concentrations of endolysin Gp105 was determined by peptidoglycan hydrolysis assay (25). Briefly, 125 μL of purified peptidoglycan was diluted with 875 μL of 60 mM MES (pH 6), 180 mM NaCl and incubated at 37°C for 30 min. Now the peptidoglycan was incubated with different concentrations (0.25 and 0.5 mg/mL) of endolysin Gp105 at 37°C and the absorbance at 600 nm were measured at regular time intervals. The peptidoglycan hydrolysis was measured by plotting the difference of absorbance at time (t) and initial absorbance (t0).

#### Colorimetric peptidoglycan hydrolysis assay

The peptidoglycan was labeled with Remazol Brilliant Blue (RBB-PG) as described earlier by Uehara et al. (26) and the peptidoglycan hydrolysis was determined based on the calorimetric assay. Briefly, to 10 μL of RBB-PG, 90 μL of PBS buffer was added and incubated at 37°C for 30 min. After incubation, 0.25 and 0.5 mg/mL of endolysin Gp105 was added to the mixture and incubated up to 6 hrs (hydrolysis period). The reaction was quenched by adding 100 μL of absolute ethanol after every one hour or hydrolysis period. The release of RBB dye was measured by pelleting the peptidoglycan at 90,000 × g at 25°C for 30 min and the absorbance (A_595nm_) was measured. The graph was plotted using the absorbance measured after 1 and 4 hour when the RBB-labeled peptidoglycan was incubated with buffer and endolysin Gp105.

#### Antibacterial assay

Anti-bacterial assay was performed as described previously (27). This assay was performed in the presence or the absence of outer membrane permeabilizers, i.e. EDTA. Briefly, the bacteria (*E. cloacae*) were grown in LB broth at 37°C to reach log-phase culture. The bacterial cells were collected by centrifugation at 6000 × g for 10 min and washed once with sterile distilled water. For EDTA treatment, the bacterial pellet was resuspended in LB broth and EDTA was added 2 mM, and incubated for 5 min at 37°C. After incubation, the cells were centrifuged at 6000 × g for 10 min and washed twice with sterile distilled water. Then, the bacterial pellet was resuspended in LB broth and used for further experiments, with controls that were not treated with EDTA but otherwise processed identically. Well-diffusion assay was performed in LB agar plates. Here, 200 μL of up to 0.8 mg/mL of endolysin was added into mechanically excised holes, “punched” wells (radius: 1 mm) and incubated at 37°C for 16 hours. The inhibition zone, where no growth was observed was recorded.

A time-course analysis of antibacterial activity was performed to determine the reduction of bacterial cell numbers over time. Briefly, log-phase bacterial cells (*E. cloacae* treated with 2 mM EDTA) were incubated with the 0.25 and 5 mg/mL of endolysin Gp105 in LB media at 37°C for 24 hours. The optical density was measured at 600 nm. In parallel, a 100 μL aliquot of bacterial cells was removed at 0, 2, 4, 6, 8, 12, 16 and 24 hours, followed by determining the number of cells (CFU) by plate counting. All the experiments were repeated three times for statistical significance.

#### Determination of broad-spectrum activity

To test the broad-spectrum antibacterial activity of purified Gp105 protein, a total of 12 Gram-negative bacteria belonging to eight genera were chosen. This includes *E. coli, K. pneumoniae, E. cloacae, A. baumannii, S*. typhi, *P. aeruginosa, S. marcescens* and *Citrobacter* sp. The four Gram-positive bacteria include two each of *Enterococcus* sp. and *S. aureus*.

Minimum inhibitory concentration (MIC) of endolysin Gp105 was determined to analyze the activity. Accordingly, micro-broth dilution method (27) was followed in which the purified endolysin Gp105 was used at a concentrations ranging from 0.007 mg/mL to 0.5 mg/mL. The endolysin Gp105 activity was determined against the log-phase bacterial culture treated with EDTA (2 mM as determined from the previous experiment). Any reduction in the bacterial turbidity was noted and the minimum concentration at which no observable bacterial growth was determined as the MIC.

## Results

In this study, we identified and characterized an endolysin from the genome of a previously characterized phage myPSH1140 which infects *Enterobacter* (22). The open reading frame (ORF) Gp105 encodes for a 164 amino acid long protein, with a putative lysozyme murein hydrolase function. Such proteins facilitate the release of progeny particles from the host bacteria by degrading the peptidoglycan layer, leading to the disintegration, or lysis, of the cells. A phylogenetic analysis showed the existence of similar types of endolysins in other phage genomes such as *Enterobacter* phage CC31, *Enterobacter* phage PG7 and *Klebsiella* phage vB_KaeM_KaAlpha, with the genome of myPSH1140 exhibiting 90% and 92% similarity with *Enterobacter* phages CC31 and PG7, respectively (22) (Fig. 1A). According to InterProScan the protein contains a T4-type_lysozyme like domain (IPR001165), belonging to the Glycoside hydrolase family 24 (IPR002196). Proteins belonging to the Glycosyl hydrolase family contain catalytic glutamate and aspartic acid residues in their active site required for the cleavage of the glycosidic bond from the substrate (28). We therefore used AlphaFold2.0 to obtain a model of the structure (Fig. 1B). The predicted structure shows the position of the key catalytic residues Glu11 and Asp20 inside a clamp-like structure with the residues facing each other, allowing the enzyme to “grab” the peptidoglycan and hydrolyze it, while sliding along the polymer network.

**Figure 1.**
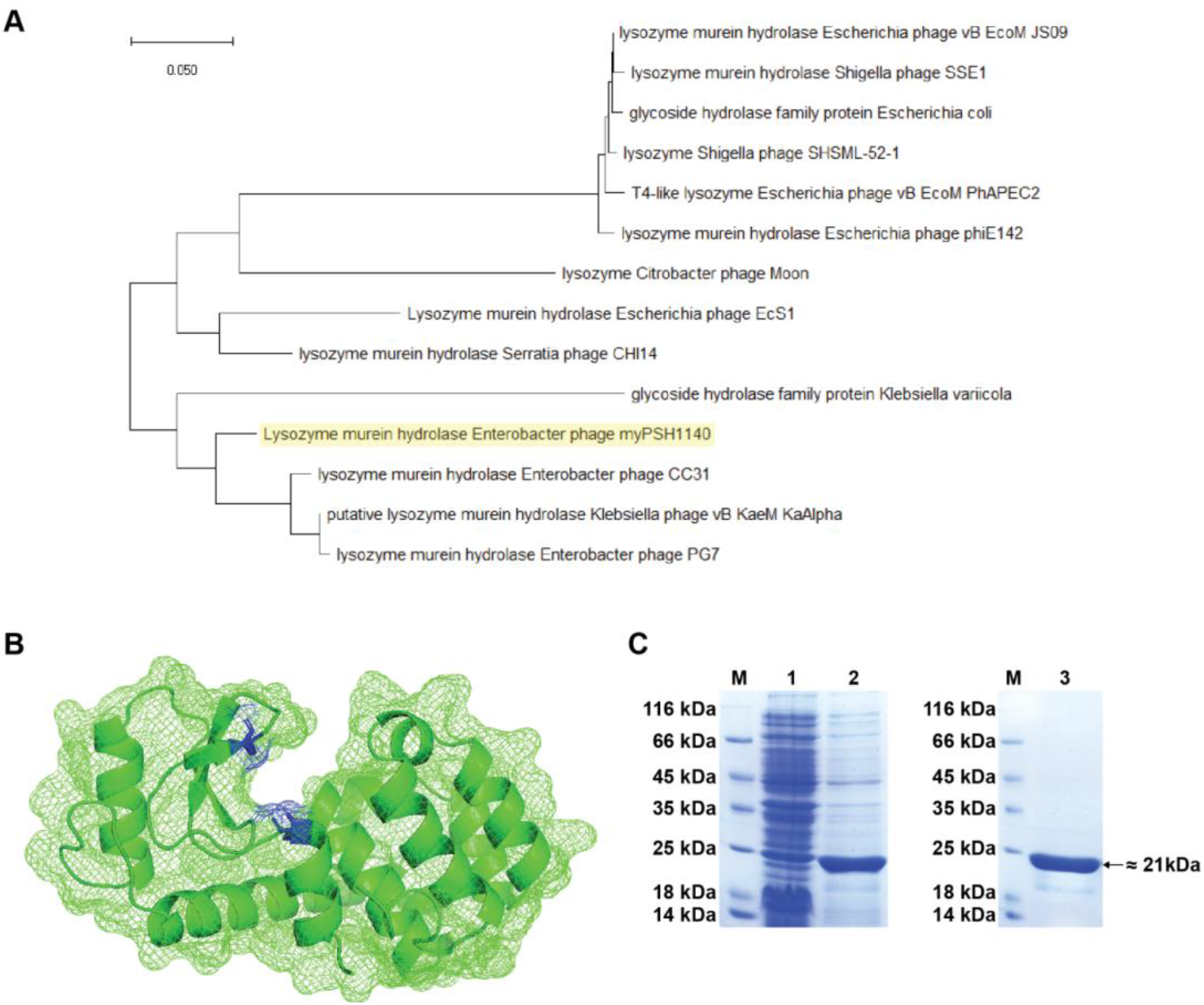
**Figure 1A. Phylogenetic analysis of Gp105 protein.** The sequences were aligned using Clustal Omega Program (https://www.ebi.ac.uk/Tools/msa/clustalo/) and a Neighbor Joining tree for conserved sites was built using MEGA software. **Figure 1B. Molecular surface diagram showing Gp105 to have tunnel like active site topology.** The proposed catalytic residues are shaded in red. **Figure 1C. SDS-PAGE analysis of Gp105 protein.** (A) Expression of Gp105 protein, Lane 1-uninduced, Lane 2-induced; Lane 3-purified. M-protein marker.

### Gp105 hydrolyses purified peptidoglycan in vitro

Since Gp105 exhibits features similar to other endolysins reported previously, and contains the key catalytic residues embedded within a predicted structure that make it likely to be able to hydrolyze peptidoglycan, we cloned the gene into an *E. coli* expression vector. The recombinant protein was produced in *E. coli* and then purified by affinity chromatography to obtain the N-terminal His-tagged protein with a molecular weight of 21 kD (Fig.1C).

To assess if the protein is functional and able to hydrolyze peptidoglycan, we purified peptidoglycan from *Enterobacter cloacae* following a protocol previously established by Santin et al. (25). Two different assays were then employed; one that makes uses of the intrinsic absorbance of the isolated material, while the second one follows the release of the Remazol Brilliant Blue dye (RBB). Using both assays, we could demonstrate that Gp105 has the ability to hydrolyze peptidoglycan from *Enterobacter*. We then tested if the enzyme exhibited a broader activity or if Gp105 is specific for the genus of *Enterobacter* (Fig. 2A). However, with peptidoglycan isolated from the Gram-positive bacterium *S. aureus* no activity was observed (Fig. 2B). Thus, endolysin Gp105 shows specificity towards its bacterial host, *E. cloacae*. Our data demonstrates that Gp105 is indeed a peptidoglycan hydrolase which has Gram-negative pathogens enzymatic specificity.

**Figure 2.**
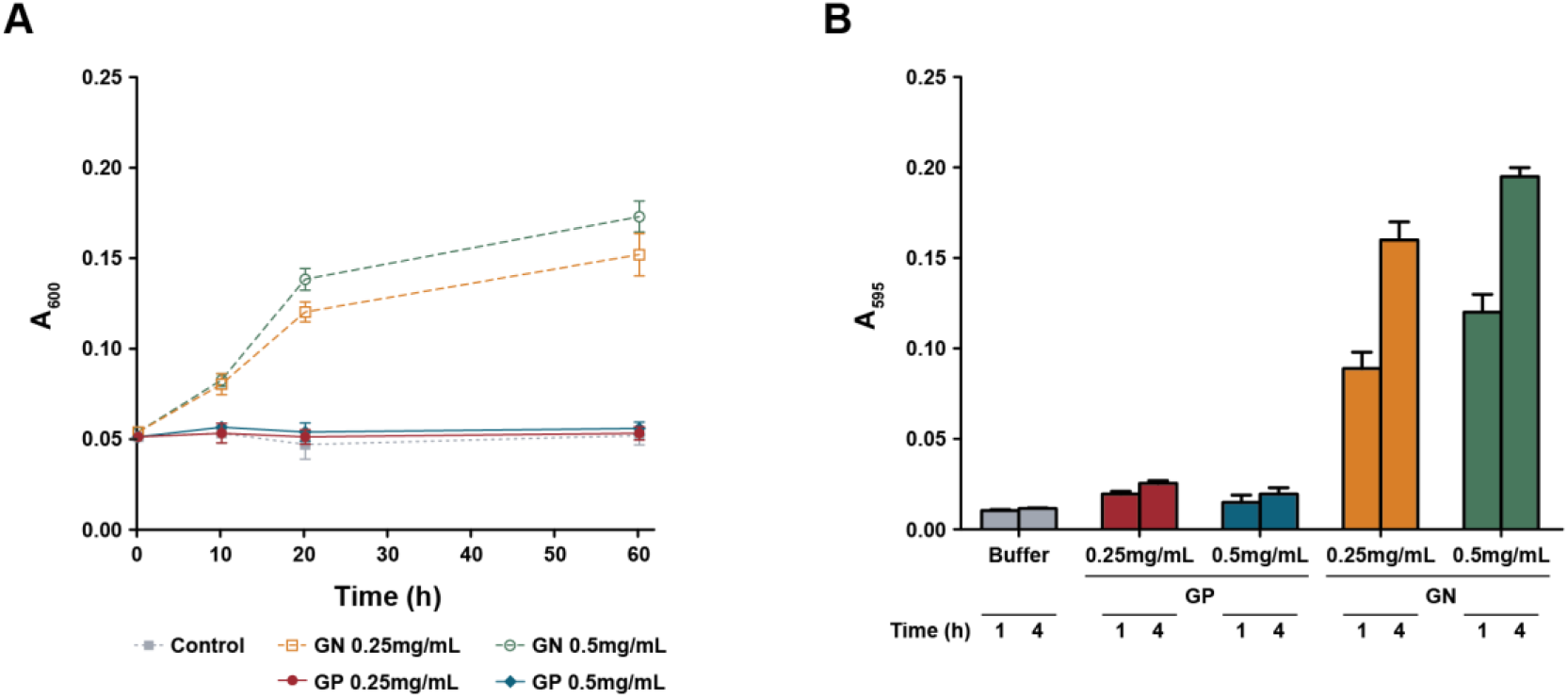
The peptidoglycan hydrolase activity of Gp105 against peptidoglycan isolated from *Enterobacter*. The peptidoglycan degradation assay was measured using absorbance (*A_600_*) at time zero (*t_0_*) and time *t* (*ΔA_600_*) **(A)** and activity measured by RBB release assay **(B)**. GP-Gram-positive bacteria, *S. aureus;* GN-Gram-negative bacteria, *E. cloacae*

### Endolysin Gp105 inhibits the growth of Gram-negative bacteria

In Gram-negative bacteria, endolysins are released into the periplasm after their synthesis, making it necessary to cross the inner membrane. Often, specialized pore-forming proteins, including holins, facilitate this process. When adding endolysins to cells from the outside, the outer membrane creates a barrier which can protect the peptidoglycan from degradation. To assess if Gp105 also has antibacterial activity, we performed two tests with 12 Gram-negative bacteria belonging to eight genera, including *E. coli*, *K. pneumoniae*, *E. cloacae*, *A. baumannii*, *S. typhi*, *P. aeruginosa, S. marcescens* and *Citrobacter* sp. in addition to four Gram-positive bacteria (two *Enterococcus* sp. isolates and two *S. aureus* strains).

In the first assay, we assessed the growth of bacteria on solid media with a reservoir of endolysin present, slowly diffusing into the agar (Table 2). No inhibition zone was observed in the case of bacterial cells that were not treated or treated with low concentrations of EDTA. Using pre-treated cells (2 mM EDTA), the strains were exposed to different concentrations of purified endolysin Gp105 at final concentrations of 0.007 mg/mL to 0.5 mg/mL. Here, clear inhibition zones were observed in the case of *Acinetobacter*, *Pseudomonas* and *Serratia* strains that were tested and to a slightly lesser extent with *Enterobacter, Klebsiella* and *Citrobacter* isolates. In contrast, we observed that neither the tested *Salmonella* nor the *Escherichia* strains showed inhibition zones in this semi-quantitative well-diffusion assay, as was also observed in the case of the Gram-positive *Enterococcus* and *Staphylococcus* isolates.

**Table 2.**
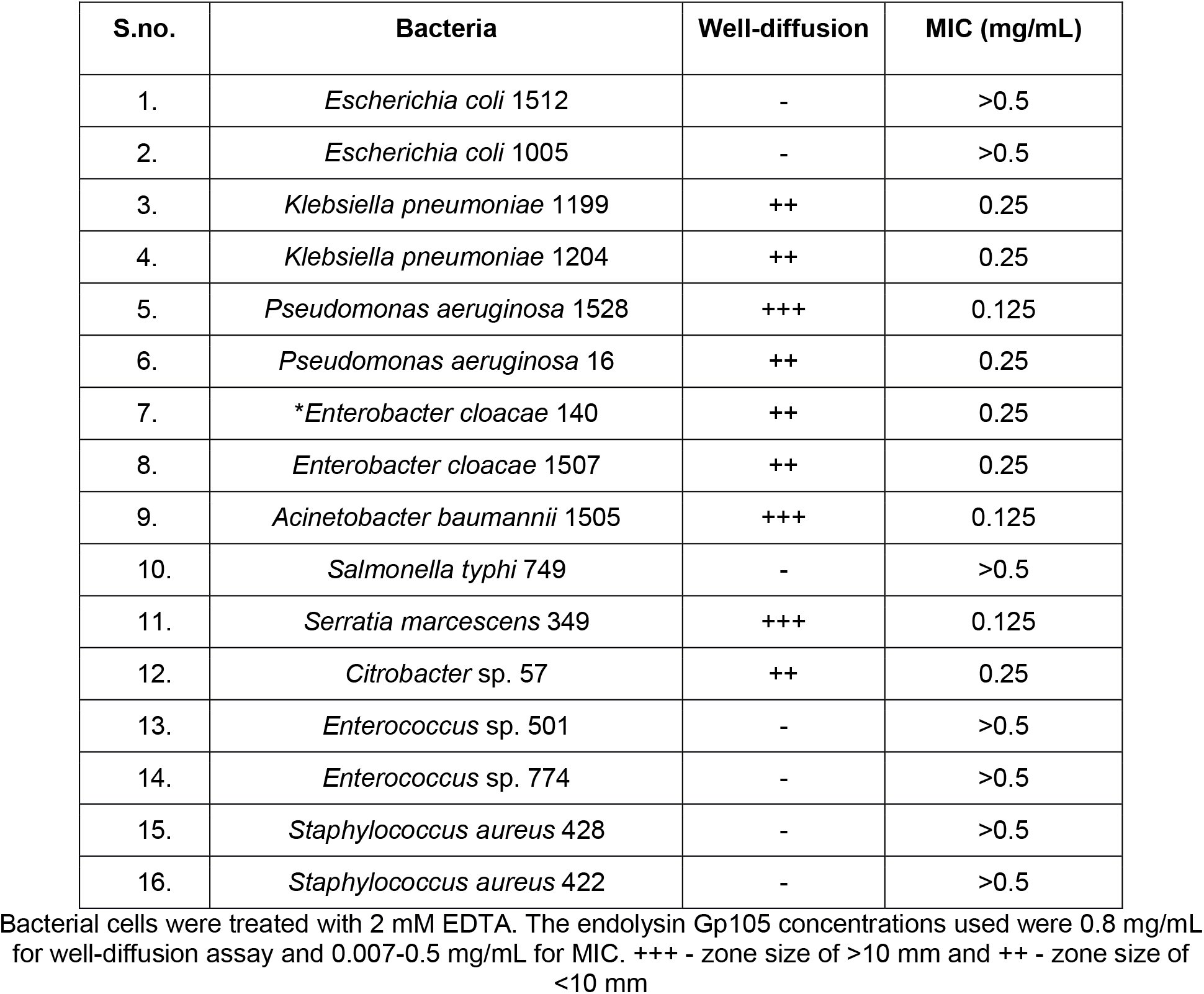
Antibacterial activity of endolysin Gp105 as observed using well-diffusion and MIC assays.

| S.no. | Bacteria | Well-diffusion | MIC (mg/mL) |
| --- | --- | --- | --- |
| 1. | *Escherichia coli* 1512 | - | >0.5 |
| 2. | *Escherichia coli* 1005 | - | >0.5 |
| 3. | *Klebsiella pneumoniae* 1199 | ++ | 0.25 |
| 4. | *Klebsiella pneumoniae* 1204 | ++ | 0.25 |
| 5. | *Pseudomonas aeruginosa* 1528 | +++ | 0.125 |
| 6. | *Pseudomonas aeruginosa* 16 | ++ | 0.25 |
| 7. | * *Enterobacter cloacae* 140 | ++ | 0.25 |
| 8. | *Enterobacter cloacae* 1507 | ++ | 0.25 |
| 9. | *Acinetobacter baumannii* 1505 | +++ | 0.125 |
| 10. | *Salmonella typhi* 749 | - | >0.5 |
| 11. | *Serratia marcescens* 349 | +++ | 0.125 |
| 12. | *Citrobacter* sp. 57 | ++ | 0.25 |
| 13. | *Enterococcus* sp. 501 | - | >0.5 |
| 14. | *Enterococcus* sp. 774 | - | >0.5 |
| 15. | *Staphylococcus aureus* 428 | - | >0.5 |
| 16. | *Staphylococcus aureus* 422 | - | >0.5 |

Bacterial cells were treated with 2 mM EDTA. The endolysin Gp105 concentrations used were 0.8 mg/mL for well-diffusion assay and 0.007-0.5 mg/mL for MIC. +++ - zone size of >10 mm and ++ - zone size of <10 mm

The second test was performed in liquid; similar to the testing of antibiotic compounds, we determined the MIC (Minimum inhibitory concentration) of the Gp105 protein (Table 2). No growth was observed at endolysin concentrations from 0.125 to 0.5 mg/mL in the case of *E. cloacae*, *K. pneumoniae, P. aeruginosa, S. marcescens, Citrobacter* sp. and *A. baumannii* (Table 2). Correlating with the results from the diffusion assay, no growth inhibition was observed in the case of *E. coli, S. typhi, E. faecalis* and *S. aureus* in the presence of the maximal concentration of the protein, 0.5 mg/ml.

Next, we determined the growth inhibition for *E. cloacae* at endolysin concentrations of 0.25 and 0.5 mg/mL. When measuring growth by determining the optical density, no significant absorbance was observed at either concentration over 24 hours compared to the control without Gp105 (Fig. 3A). When counting the CFUs at different time points, we were able to obtain live bacteria. However, over time the number did not show any major increase (Fig. 3B). Thus, the protein appears to show bacteriostatic properties, which is unusual for endolysins. A possible explanation might be that the enzyme only acts on dividing cells where it might be able to traverse the outer membrane barrier.

**Figure 3.**
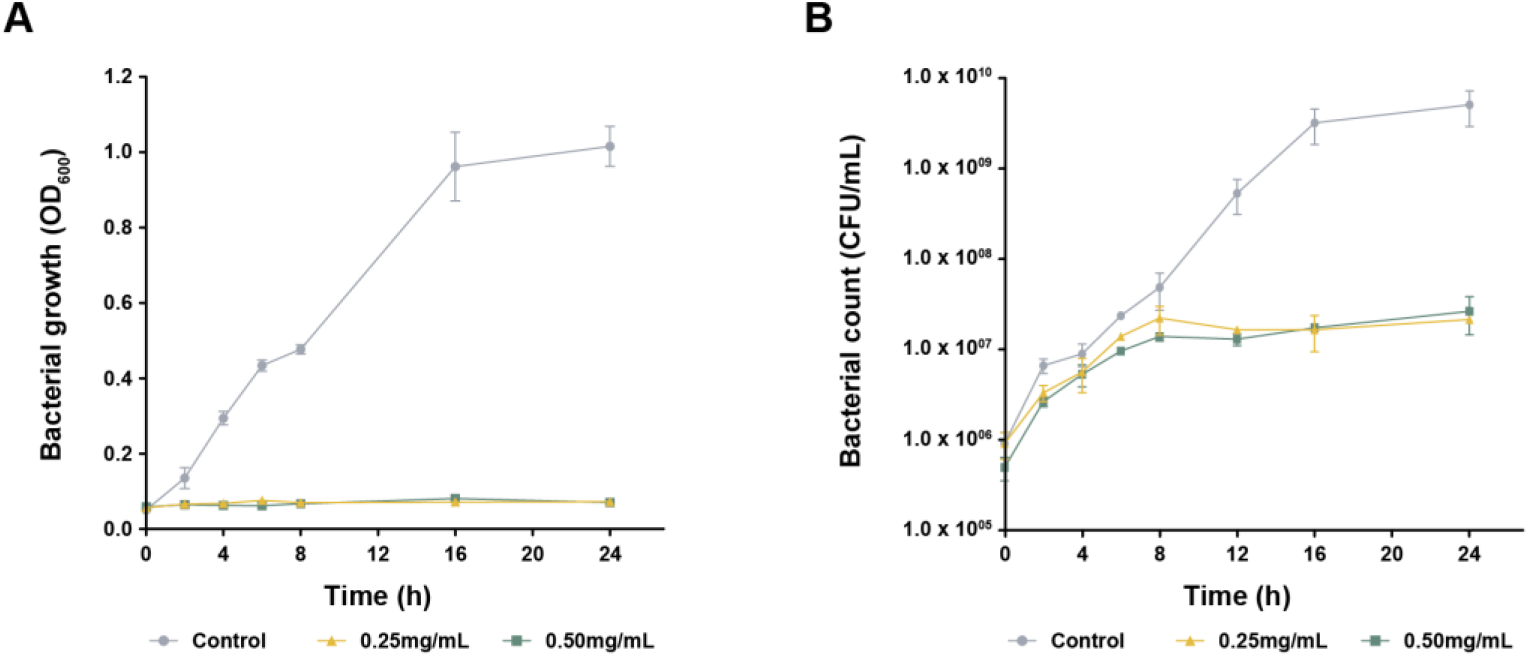
Antibacterial activity of lysozyme murein hydrolase, Gp105 against *Enterobacter cloacae*. **(A)** Optical density measurement to determine growth inhibition of *E. cloacae* pre-treated with 2 mM EDTA when exposed to 0.25 and 0.5 mg/mL of Gp105 final concentration. **(B)** The viability of cells was assessed by counting the number of colony forming units (CFUs). Data shown are standard mean values obtained from repeated experiments.

## Discussion

Antibiotic resistance is rapidly developing into a major global health pandemic, creating a requirement to identify and develop new treatment options employing chemical or biological antimicrobial compounds. Endolysins as therapeutic proteins are considered to be an effective alternative to treat bacterial infections (29). Based on the specificity of the enzyme, endolysins can be categorised into five main groups: (a) N-acetylglucosaminidases, (b) N-acetylmuramidases, which cleaves the glycan bond found on the reducing side of the MurNAc; these are also called lysozymes, (c) lytic transglycosylases, d) N-acetylmuramoyl-L-alanine and (e) peptidases (30).

The endolysin we investigated is from an *Enterobacter* phage which is underrepresented in published research. To the best of our knowledge, this is the first report of an endolysin from a phage infecting *Enterobacter* sp. In this study, the purified protein Gp105 with a predicted lysozyme murein hydrolase function from *Enterobacter* phage myPSH1140 was found to have broad-spectrum antibacterial activity against Gram-negative bacteria. The hydrolase activity was also confirmed by using peptidoglycans isolated from *E. cloacae*.

Gp105 is a T4-type_lysozyme domain containing protein which belongs to the Glycosyl hydrolase family. The T4 lysozyme hydrolyses the 1,4-beta linkages between N-acetyl-D-glucosamine and N-acetylmuramic acid in peptidoglycan heteropolymers of bacterial cells (28). Further studies are required to determine if the mechanism behind the enzymatic activity of Gp105 is the same as that of T4 lysozyme.

The recombinant expression of Gp105 in *E. coli* allowed the high-yield production of a functional protein with yields of approximately 1-2 mg of purified protein per mL of culture. The high yield expression is possibly due to the inactivity of the protein towards *E. coli*, as also shown in the results of the well-diffusion assay. The observed molecular weight of 21 kDa is similar to that of other endolysins previously reported which exhibit sizes between 16 and 25 kDa (31,32).

Especially promising is the fact that Gp105 shows broad-spectrum activity against at least six Gram-negative pathogens. However, as with all other endolysins acting on Gram-negative bacteria, cells were treated with EDTA which binds and removes divalent ions that stabilize the surface structure of cells, thus resulting in a partially destabilized outer membrane. Other studies reported have also employs such compounds to study the activity of the enzymes against Gram-negative bacteria (33). While the use of chemicals such as EDTA is medically not possible, the key aspect of studies such as ours and previously reported research is the activity of the enzyme per se, albeit in concert with EDTA. Adding amino acids such as Lysin or Arginine which are opposite charged to the cell envelope or creating fusion proteins with cell-membrane penetrating peptides, might allow the use of such synthetic enzymes without EDTA in the future.

## Conclusion

In our study we characterized a lysozyme murine hydrolase, Gp105, from *Enterobacter* phage myPSH1140 and analysed its structure and antibacterial activity *in silico, in vitro* and *in vivo*. Gp105 is a T4-type lysozyme domain containing protein which belongs to the Glycosyl hydrolase family. The identified peptidoglycan hydrolase gene was cloned, expressed and purified to explore its ability to inactivate Gram-negative bacteria. The broad-spectrum activity of Gp105 could be exploited to inhibit or kill pathogenic bacteria in therapeutic applications.

## Acknowledgments

The authors would like to thank the Vellore Institute of Technology for providing VIT SEED grant. This research work was supported by Zhejiang Province Post-Doctoral Research Fund received by PM.

